# A New Framework for MR Diffusion Tensor Distribution

**DOI:** 10.1101/2020.05.01.071118

**Authors:** Kulam Najmudeen Magdoom, Sinisa Pajevic, Gasbarra Dario, Peter J. Basser

## Abstract

The ability to characterize heterogeneous and anisotropic water diffusion processes within macroscopic MRI voxels non-invasively and *in vivo* is a *desideratum* in biology, neuroscience, and medicine. While an MRI voxel may contain approximately a microliter of tissue, our goal is to examine intravoxel diffusion processes on the order of picoliters. Here we propose a new theoretical framework and experimental design to describe and measure such intravoxel structural heterogeneity and anisotropy. We assume that a constrained normal tensor-variate distribution (CNTVD) describes the variability of positive definite diffusion tensors within a voxel which extends its applicability to a wide range of b-values unlike existing models. We use a Monte Carlo scheme to synthesize realistic numerical diffusion tensor distribution (DTD) phantoms and invert the MR signal. We show that the signal inversion is well-posed and estimate the CNTVD parameters parsimoniously by exploiting the different symmetries of the mean and covariance tensors of CNTVD. The robustness of the estimation pipeline is assessed by adding noise to calculated MR signals and compared with the ground truth. A family of invariant parameters which characterize microscopic shape, size and orientation heterogeneity within a voxel are also presented.

## Introduction

Diffusion tensor imaging (DTI) is a widely accepted MR imaging method used by clinicians and neuroscientists to assess microstructure and organization of brain and other soft tissues [1, 2]. DTI consists of measuring a single (mean) diffusion tensor within each voxel of an imaging volume and allows characterization of brain tissue microstructure, architecture, organization, and anatomical connectivity. Applications have been widespread in brain and whole body clinical imaging [3, 4].

However, DTI only provides an estimate of a single mean diffusion tensor within each voxel, which is typically on the order of one cubic millimeter, representing a microliter of tissue. But, at microscopic, mesoscopic, and even macromolecular length scales the brain and other tissues contain a myriad of cell sizes and types, fine processes, extracellular matrix (ECM) and intracellular components with complex microstructural and architectural organizations [5]. Histological analysis often reveal distinct anisotropic domains within the same voxel, such as multiple white matter pathways which may interdigitate, kiss, cross, merge, diverge, etc. A goal of “in vivo MRI histology” or “microstructure imaging” has been to achieve subvoxel resolution and extract useful information about various salient water pools to provide a more nuanced and comprehensive assessment of the complex microscopic tissue milieu. Such information could be invaluable in understanding of normal and abnormal development, neuropathological changes, and brain structure-function relationships, to name a few.

The fundamental but coarse-scale description of diffusion anisotropy within a voxel, resulting from an anisotropic Gaussian diffusion process, is via DTI [1, 2]. Several approaches were later introduced to capture both the heterogeneous and anisotropic character of diffusion within a MRI voxel. Tuch et al. introduced high angular resolution diffusion imaging (HARDI) where the MR signal is acquired in more than the six directions, the minimum number required by DTI, and the diffusivity was expressed as an angular function to resolve multiple fibers within a voxel [6]. Frank refined this approach by expressing the measured diffusivity in each direction using spherical harmonics [7]. Ozarslan and Mareci introduced generalized DTI by representing the HARDI signal using higher order tensors (HOT) to overcome ambiguities with DTI model [8]. Liu et al. and later Jensen et al. introduced the kurtosis tensor to account for the MR signal not captured by the DTI model which were assumed to arise from non-Gaussian diffusion of water in brain tissue [9, 10].

An important advance in the field was made by Jian et al. where the neural tissue is modeled to be composed of microscopic Gaussian diffusion compartments described by the joint probability distribution for the six elements of the diffusion tensor (i.e. diffusion tensor distribution or DTD) which in general is unknown [11]. Obtaining the 6D DTD non-parametrically involves taking the inverse Laplace transform (ILT) of the MR signal, which is a well-known ill-posed inverse problem [12]. To overcome the ill-posedness, Jian et al. assumed a discrete mixture of Wishart distribution for the DTD and showed it could resolve crossing fibers. Leow et al. reduced the DTD from 6D to 4D by assuming cylindrical symmetry and numerically computed it for the brain tissue [13]. However, the aforementioned studies used rank-1 b-matrices resulting from their use of single pulsed-field gradient (sPFG) to obtain diffusion weighting, which limit the detection of microscopic anisotropic domains within a voxel [14, 15] which have been shown to exist in the brain and spinal cord [16, 17, 18]. The kurtosis model is also limited by the unphysical increase in the diffusion weighted MR signal with b-value [19]. This limits the amount of microstructure information that could be gleaned from it given that DTI successfully accounts for most of the signal (approximately 90 percent) at low b-values [20]. The critical b-value is in general unknown since it is dependent on the underlying kurtosis [19].

Recently, Topgaard obtained cylindrically symmetric 4D DTD in a phantom using higher rank b-matrix measurements with Monte Carlo (MC) signal inversion [21, 22]. A key aspect of their work is to classify the underlying microstructure in terms of size, shape and orientation heterogeneity. The method however is time consuming, requiring almost five hundred volumes. Using spherical encoding, Avram et al. obtained the distribution for the trace of the diffusion tensors in the brain tissue from ILT of the data with regularization and showed it to be Gaussian [23]. Callaghan introduced the cumulant expansion of MR signal for scalar diffusivity whereby the MR signal is approximated in terms of the moments of the DTD which are related to underlying microstructure [12]. The advantage of this method is that it does not assume a particular DTD but the expansion has a limited radius of convergence causing unphysical signal increase with b-value similar to the kurtosis model [19]. The cumulant description is also not compact in higher dimensions since the number of unknown parameters increases exponentially with increasing order of the HOT required to describe these moments. Westin et al. extended the cumulant approach to 6D DTD with higher rank b-matrices and applied on the neural tissue for low diffusion weighting (b < 2.5 ms/*μ*m^2^) where the expansion is assumed to be valid [24]. The cumulant expansion was performed to include the first two moments of the DTD with the second moment (i.e., the covariance) assumed to be isotropic which often is not rich enough to describe the microstructural tissue complexity.

In this work, a novel paradigm is introduced to obtain the 6D-DTD within a voxel which overcomes the limitations discussed above. The DTD is assumed to be a tensor-variate normal distribution [25] but constrained to lie on the manifold of positive semi-definite diffusion tensors (CNTVD), parameterized by second-order mean and fourth-order covariance tensors. We show that the CNTVD has maximum entropy property which makes it a natural choice for DTD, and allows us to admit larger b-values to obtain microstructural information not captured by the DTI, cumulant and kurtosis models. We also show that rank-1 and rank-2 b-matrix measurements are sufficient for estimating the mean and covariance tensors, and a compressed sensing (CS) based experimental design is introduced to perform the MR measurements efficiently. We also introduce a new MC scheme to synthesize realistic voxel wise 6D DTDs, and invert the signal which we show as a well-posed problem. A parsimonious model selection framework exploiting the symmetries of the mean [26] and covariance tensors [27] is introduced to estimate the CNTVD parameters. The estimated parameters are compared with the ground truth to assess their accuracy and precision. The micro-orientation distribution function (*μ*ODF), micro-fractional anisotropy (*μ*FA) and a family of invariant stains are introduced to characterize the intravoxel size, shape and orientation heterogeneities.

## Methods

### MR signal model

The MR signal from an ensemble of diffusion tensors distributed according to *p*(**D**) is given by [11],

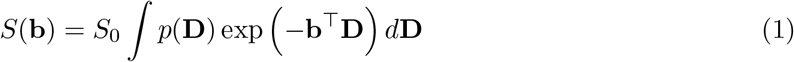

where *S*_0_ is the signal without diffusion weighing, **D** = (*D*_*xx*_, *D*_*yy*_, *D*_*zz*_, *D*_*xy*_, *D*_*xz*_, *D*_*yz*_)^⊤^ is a vector of the independent components of the second order symmetric diffusion tensor, and **b** = (*b*_*xx*_, *b*_*yy*_, *b*_*zz*_, 2*b*_*xy*_, 2*b*_*xz*_, 2*b*_*yz*_)^⊤^ is a vector of the independent components of the 3 × 3 symmetric diffusion weighting b-matrix [1]. For a normal distribution of tensors constrained in the manifold of positive definite tensors, 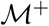, *p*(**D**) is given by,

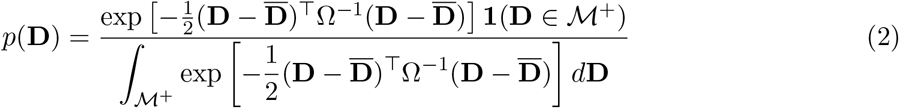

where **1** is the indicator function and 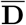 is second order mean diffusion tensor expressed as a 6 × 1 vector, and Ω is the fourth order covariance tensor, C, expressed as a 6 × 6 matrix [27] according to the following mappings,

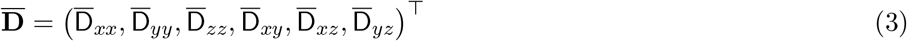

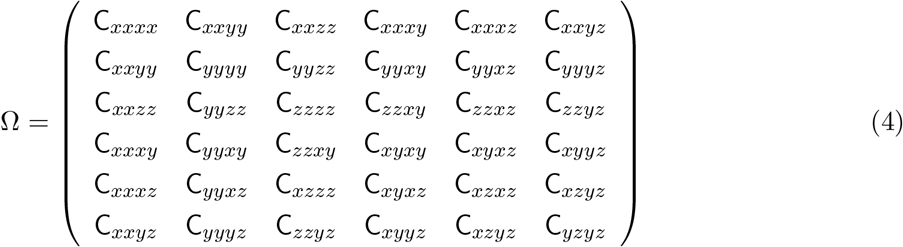

The signal equation is approximated using MC integration with samples, **D**_*i*_, drawn from the CNTVD as shown below,

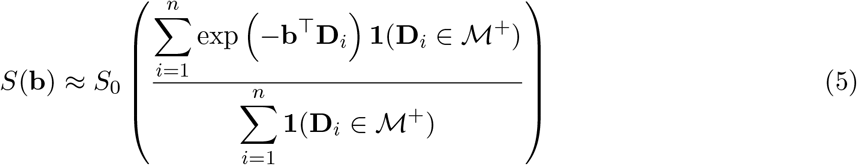

where *n* is the number of MC samples, set to 200,000 in our simulations. The choice of CNTVD for DTD is motivated by its resemblance to that measured in the brain tissue for orientationally averaged diffusivity [23], and a few of its properties which we prove in Section 1 in the supplementary material. Given that DTD is generally unknown in a voxel, we show that CNTVD is the natural choice since it is the least informative distribution (or maximum entropy) among all probability densities supported by a constraint (e.g., positive definiteness of diffusion tensors) with a given first and second moments. This maximum entropy property of CNTVD avoids adding information that are not present [28]. We also prove that the moments of CNTVD are unique for a given DTD which renders the data inversion problem well-posed.

For comparison, MR signal from the cumulant expansion and kurtosis model were also computed for the given mean and covariance. The kurtosis tensor, K_*ijkl*_, subsumed by the covariance tensor, is obtained by imposing additional symmetries on the covariance tensor resulting in the following identity (Section 2 in the supplementary material),

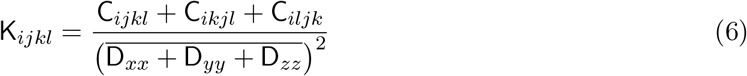

### DTD synthesis

The individual micro-diffusion tensors (i.e., MC samples) in the signal equation (Equation (5)) are synthesized from random samples drawn from 6D normal multivariate distribution with mean and covariance that define the desired DTD. The 6D samples are reformulated into 3 × 3 second order tensors using the equivalence between the two formulations shown by Basser and Pajevic [27]. A positive definiteness filter is applied on the drawn samples to satisfy the physical constraint on the diffusion tensor. In addition to obtaining the signal, the synthesized micro-diffusion tensors also demonstrate the richness of CNTVD.

Since the mean and covariance of a DTD is in general unknown *a priori*, it was calculated empirically from a sample of micro-diffusion tensors generated by randomly varying their eigenvalues and/or eigenvectors. For example, samples for an isotropic emulsion are generated from individual isotropic micro-diffusion tensors such that their trace follows a univariate normal distribution.

### Experimental design

A b-matrix of rank-*m*, b_*m*_, (with *m* ∈ {1, 2, 3}) is generated from diffusion encoding gradient vectors, **q**_*i*_, as shown below,

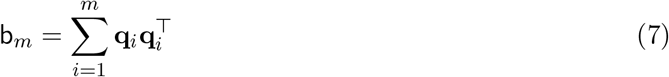

It is impractical to span the entire space of b-matrices of all three ranks. However, using tensor algebra we prove that a combination of rank-1 and rank-2 b-matrices are sufficient to span the entire space of fourth-order covariance tensors (Section 3 in the supplementary material). Further data reduction is achieved using CS which has been employed in several multi-dimensional MR studies exploiting the sparsity in *q*-space [29, 30, 31]. A combination of rank-1 and rank-2 b-matrices were obtained for DTD using double pulsed field gradient (dPFG) or double diffusion encoding measurements [14, 15, 16]. The two gradient vectors in dPFG were randomly oriented uniformly over a sphere with their lengths uniformly distributed to obtain a range of b-values of interest. The resulting collection of b-matrices have varying orientation, size and shape, an example of which is shown in Figure 1.

**Figure 1:**
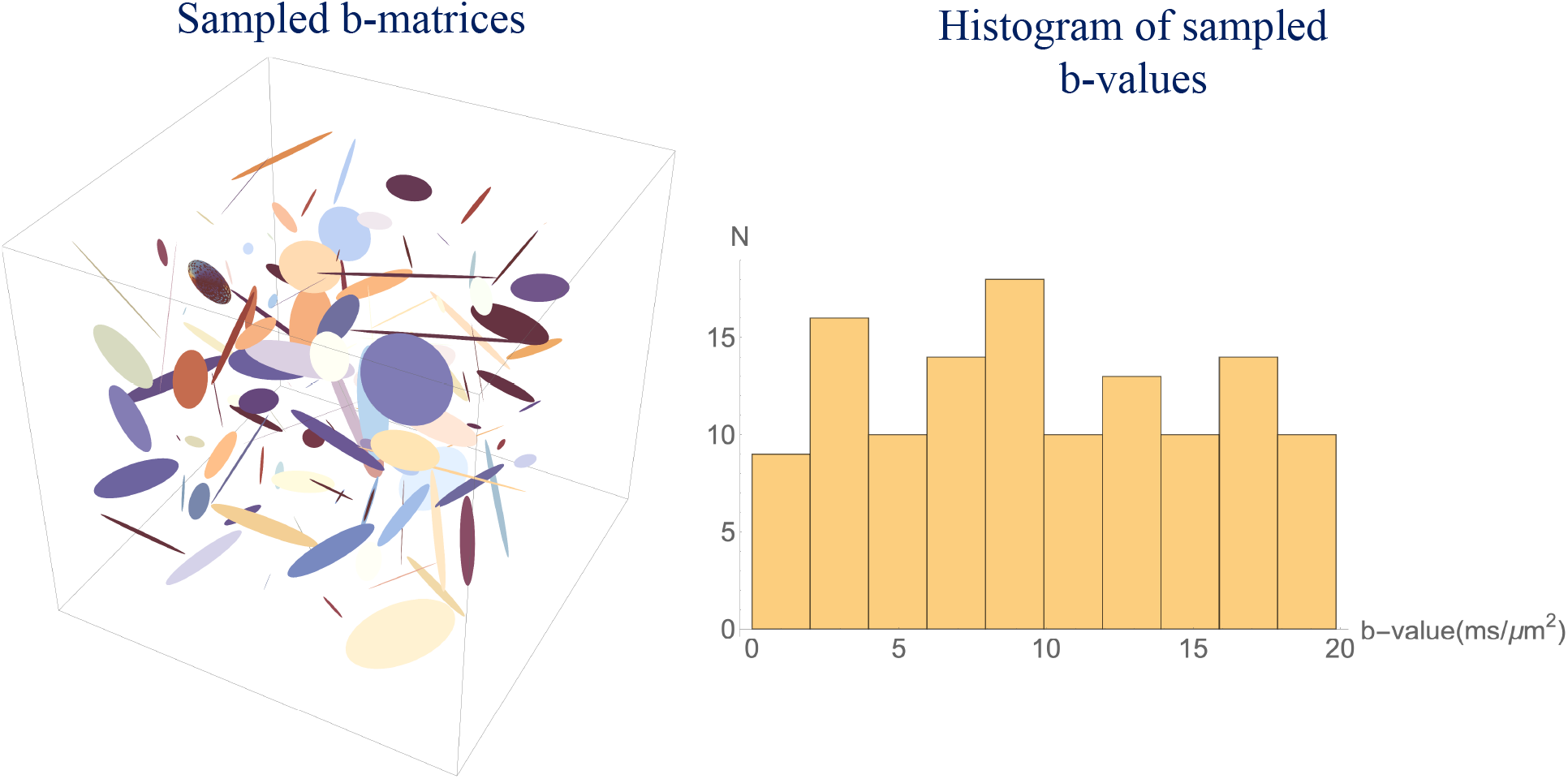
Experimental design for DTD estimation. (Left) An example of the sampled rank 1,2 b-matrices shown as ellipses generated using randomly oriented and sized diffusion gradient vectors. (Right) Histogram of trace of b-matrix showing the uniform sampling in size.

### Parameter estimation

For a given DTD, the MR signal is calculated for the designed b-matrices (*N* = 125) using Equation (5). Gaussian noise is added on the real and imaginary channels of the MR signal, if necessary, before inverting the signal to obtain the CNTVD parameters characterizing the DTD. The number of unknowns for the most general DTD is 28 (6 for mean, 21 for the covariance and 1 for *S*_0_), which makes the signal inversion susceptible to over-fitting [32]. A family of physically plausible nested models for the mean and covariance are then used to balance the bias and variance in the estimated parameters.

The mean tensor including *S*_0_ is classified into the following four submodels: 1) 1-parameter noise model (i.e., only *S*_0_), 2) 2-parameter isotropic model, 3) 5-parameter prolate/oblate models and 4) 7-parameter general anisotropic model [26]. The equivalence between the 6×6 covariance matrix and the fourth order tensor [27] is used to classify the covariance tensor into the following eight submodels exploiting symmetries of the fourth order tensor well known in the elasticity literature: 1) 2-parameter isotropic model, 2) 3-parameter cubic model, 3) 5-parameter hexagonal model, 4) 6-parameter trigonal model, 5) 7-parameter tetragonal model, 6) 9-parameter orthorhombic model, 7) 13-parameter monoclinic model, and 8) 21-parameter triclinic model (see page 140-141 in [33]).

For a given submodel, the MR signal is fit to Equation (5) using simplex-type numerical optimization algorithm implemented in Python to estimate the unknown parameters. The Bayesian information criteria (BIC) is used for selecting models with greatest parsimony, (i.e., which provide an optimal trade-off between the goodness-of-fit and the number of free parameters) [32]. The model with the fewest parameters is selected unless the difference in its BIC with a more complex model is greater than two [34]. The appropriate mean tensor model is first selected from the data with the covariance set to zero to account for uniform DTDs. The selected mean tensor model is then augmented with each of the eight covariance models to find the one which parsimoniously describes the data.

### Microstructural stains and glyphs

The estimated CNTVD parameters are used to delineate several microstructural features within the voxel. The micro-diffusion tensors within the voxel are simulated by drawing samples from CNTVD with the estimated parameters. The mean tensor is visualized as an ellipsoid [2], while the fourth order covariance tensor is visualized as a 3D glyph by projecting the tensor onto a 3D sphere using the tensor contraction, *r_i_r_j_r_k_r_l_*C_*ijkl*_, where **r**(*θ*, *ϕ*) = (sin *θ* cos *ϕ*, sin *θ* sin *ϕ*, cos *θ*) is the unit vector on a sphere [27, 35]. The radius of the glyph along a given orientation is set to the projected value of the covariance in that direction.

Given the mean and covariance of CNTVD, microscopic quantities such as micro-fractional anisotropy (*μ*FA), *μ*ODF and micro-entropy (*S*_*D*_) are computed using the following relation,

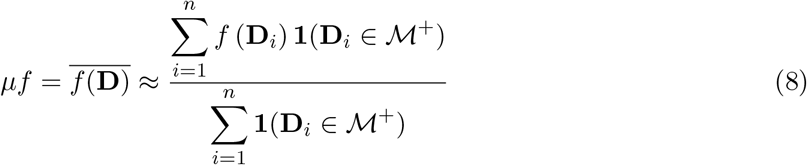

where *f* is the function of interest such as FA, ODF or entropy (i.e.− log *p*(**D**)). Leow et al. introduced the *μ*ODF definition but without using the Jacobian for proper radial integration of the propagator [13]. Aganj et al. obtained the ODF for the DTI model with the correct Jacobian for integration [36] which was used in our calculation. In contrast to the scalar *μ*FA quantity, the macro- and micro-ODFs within a voxel are displayed as 3D glyphs where the radius of the glyph along a given orientation is equal to the value of the ODF along that direction.

The size heterogeneity within a voxel is expressed as the coefficient of variation for mean ADC (mADC) or average trace of the individual micro diffusion tensors. Given a multivariate normal random variable, **D**, it is well known that the distribution of a scalar projection of **D**, **C**^⊤^**D**, where **C** is a 1×6 random constant vector, has mean, **C**^⊤^**D**, and variance, **C**^⊤^Ω**C** [37]. Using this identity, variance of mADC is obtained by choosing **C** equal to 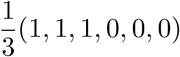. This results in the mADC variance equal to the arithmetic average of upper 3 × 3 block matrix appearing in Ω from which the expression for coefficient of variation for size, CV_size_, is given below,

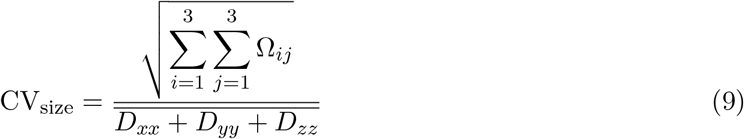

Given the shape of an ellipsoid is uniquely determined by the ratio of its eigenvalues [38], the shape heterogeneity within a voxel is expressed as the coefficient of variation of the eigenvalue ratios of micro-diffusion tensors, CV_shape_, as given below,

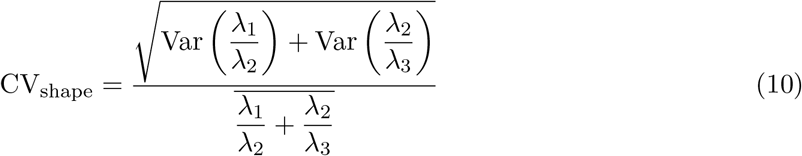

where *λ*_1_ > *λ*_2_ > *λ*_3_ are the individual eigenvalues of the micro-diffusion tensor and Var is the variance. The orientation heterogeneity within a voxel is expressed as the gain in ODF entropy from averaging, Δ*S*_ODF_, as given below,

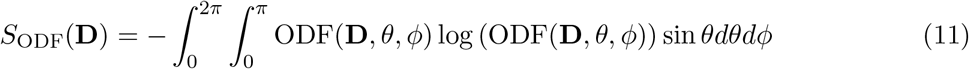

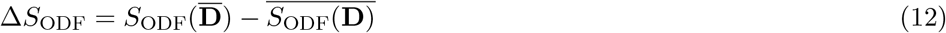

where *S*_ODF_(**D**) is the ODF entropy for the diffusion tensor, **D**.

## Results

The DTI model is unable to distinguish among various DTDs that might adequately describe diffusion processes in gray and white matter. Some of those microstructural templates can be generated using CNTVD as shown in Figures 2 and 3. The figure also shows the ability of CNTVD to remove this ambiguity by assigning distinct covariance matrices corresponding to each of these DTDs even though their mean diffusion tensors are identical (also shown in the figures). It also shows how a single covariance matrix could describe a mixture of size, shape and orientation heterogeneities that maybe present within a MRI voxel.

**Figure 2:**
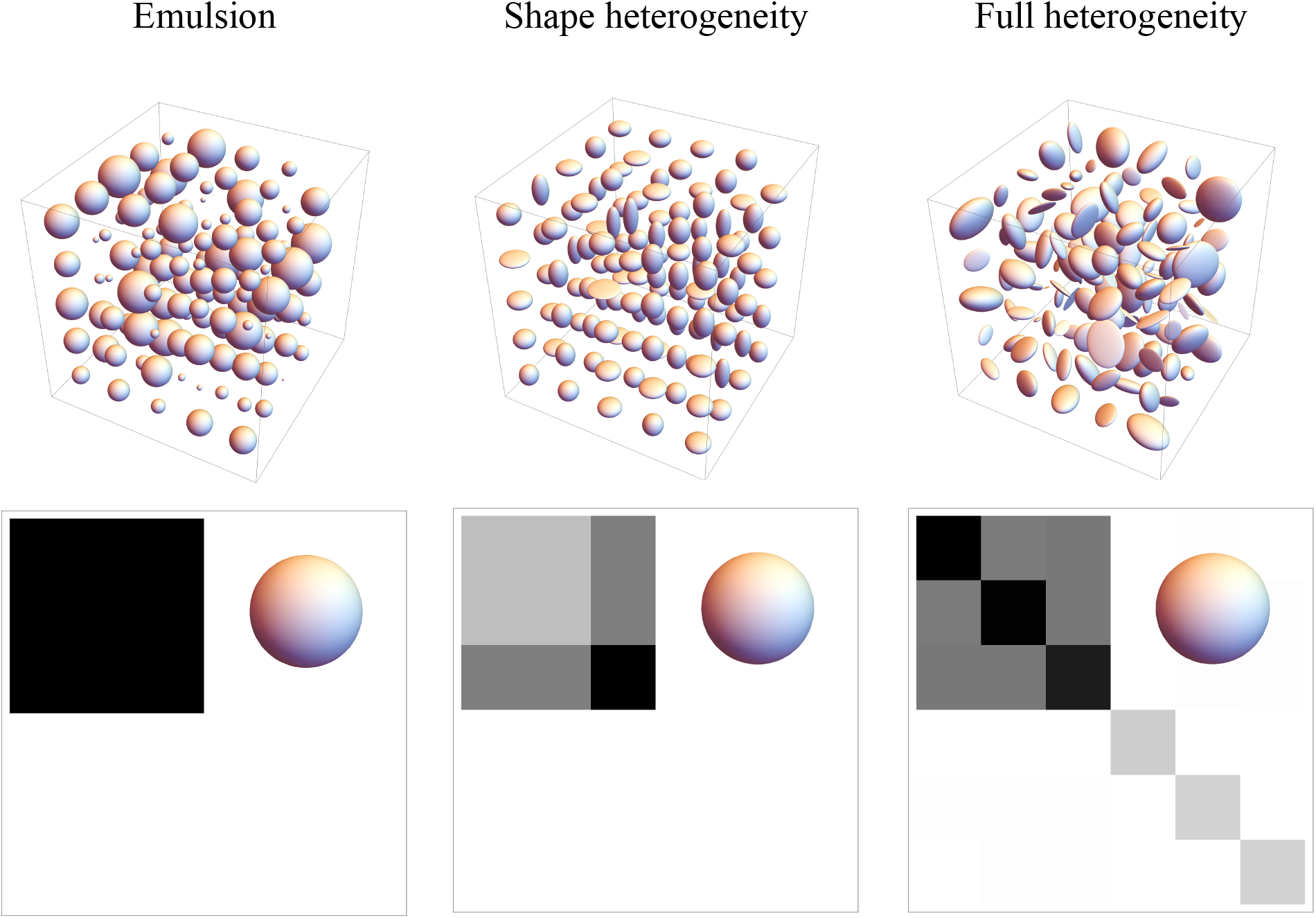
Macroscopically isotropic and microscopically heterogeneous DTDs generated from CNTVD describing gray matter. The structure of the 6 × 6 covariance matrix along with the mean tensor (inset) is shown in the bottom row. (Left column) Emulsion type DTD with size heterogeneity, (Middle column) Shape heterogeneous DTD with a mixture of prolate, oblate and spherical diffusion tensors, (Right column) Fully heterogeneous DTD with randomly oriented tensors of varying shapes and sizes.

**Figure 3:**
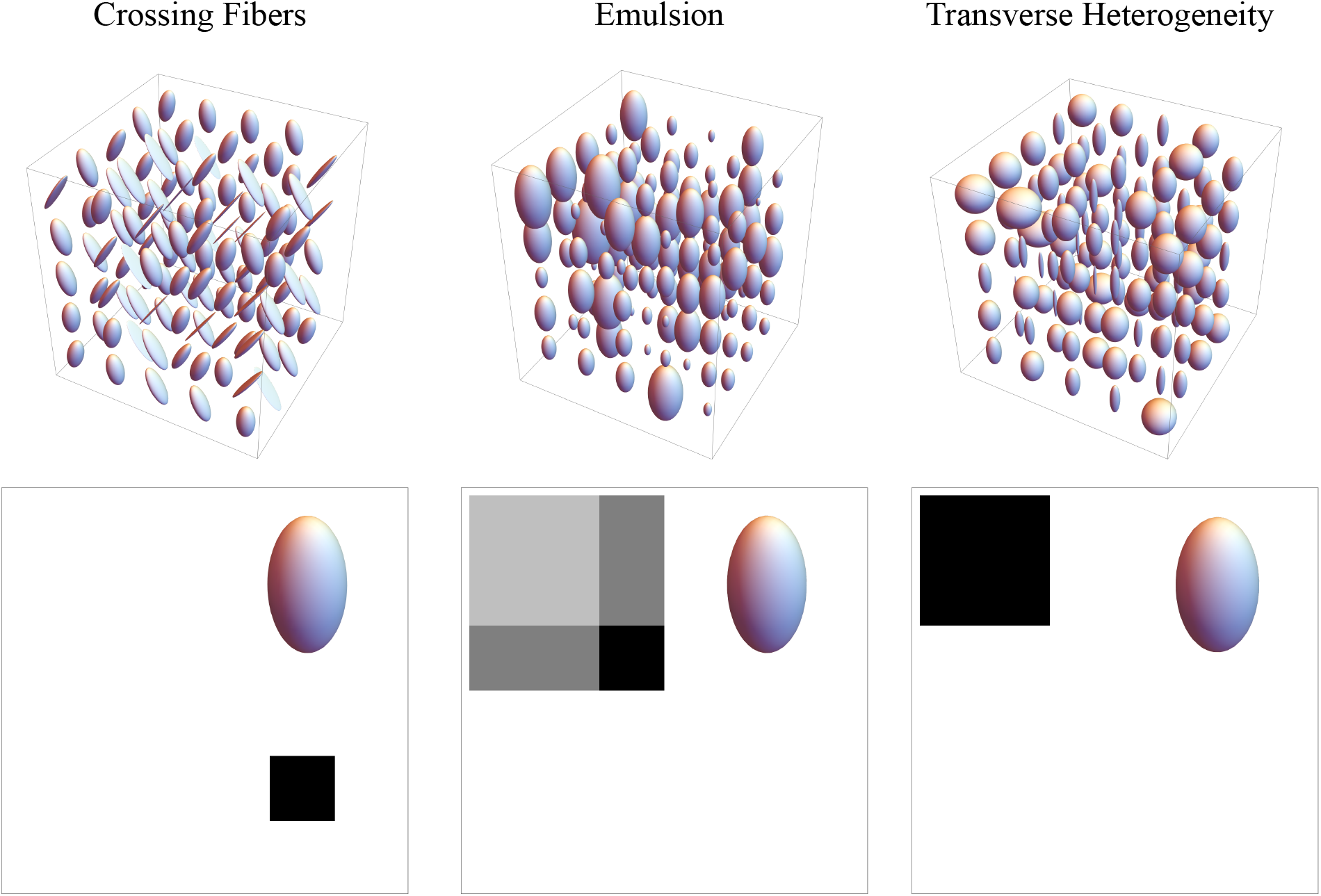
Macroscopically anisotropic and microscopically heterogeneous DTDs generated from CNTVD describing the white matter. The structure of the 6 × 6 covariance matrix along with the mean tensor (inset) is shown in the bottom row. (Left column) Crossing fiber DTD with 90-degree angle between the fibers, (Middle column) Emulsion type DTD with size heterogeneity, (Right column) DTD with random transverse eigenvalues characterizing a bundle of white matter fibers of varying diameters.

A gray matter voxel is modeled as macroscopically isotropic but microscopically composed of diffusion tensors of varying size, shape (i.e., prolate, oblate and sphere) and orientation as shown in Figure 2. The size and fully heterogeneous (i.e., size, shape and orientation heterogeneous) DTDs exhibited an isotropic covariance structure characterized by two constants, *λ* and *μ*, similar to the bulk and shear modulus of the isotropic elasticity tensor [25, 27]. The size heterogeneous DTD had *λ* ≠ 0 and *μ* = 0 while fully heterogeneous DTD had *λ* ≠ 0 and *μ* ≠ 0. The shape heterogeneous DTD exhibits a hexagonal covariance structure.

A white matter voxel is modeled as macroscopically anisotropic and microscopically composed of distinct anisotropic subdomains shown in Figure 3. The three DTDs considered are two fibers crossing at 90 degrees, anisotropic emulsion, and an axon bundle consisting of varying diameter fibers modeled by randomly varying the transverse eigenvalues together but fixing the longitudinal eigenvalue of the micro-diffusion tensors. The crossing fiber exhibited an orthorhombic covariance structure while the other two DTDs exhibited a hexagonal structure.

The MR signal for some of the aforementioned DTDs obtained using the proposed model is compared with DTI, kurtosis models and cumulant approximation in Figure 4. Since *p*(**D**) for these DTDs is the CNTVD, the proposed model is the ground truth for the MR signal. It can be observed that DTI fails to predict the signal at high b-values despite its monotonic decay with b-value. The kurtosis model and cumulant approximation however show an unphysical signal increase with b-value which is not observed experimentally. In addition, it is seen that they begin to deviate from the ground truth at very low b-values, well before the minimum in the signal decay curve occurs; thus cumulant approximation and kurtosis model are only valid for very low b-values.

**Figure 4:**
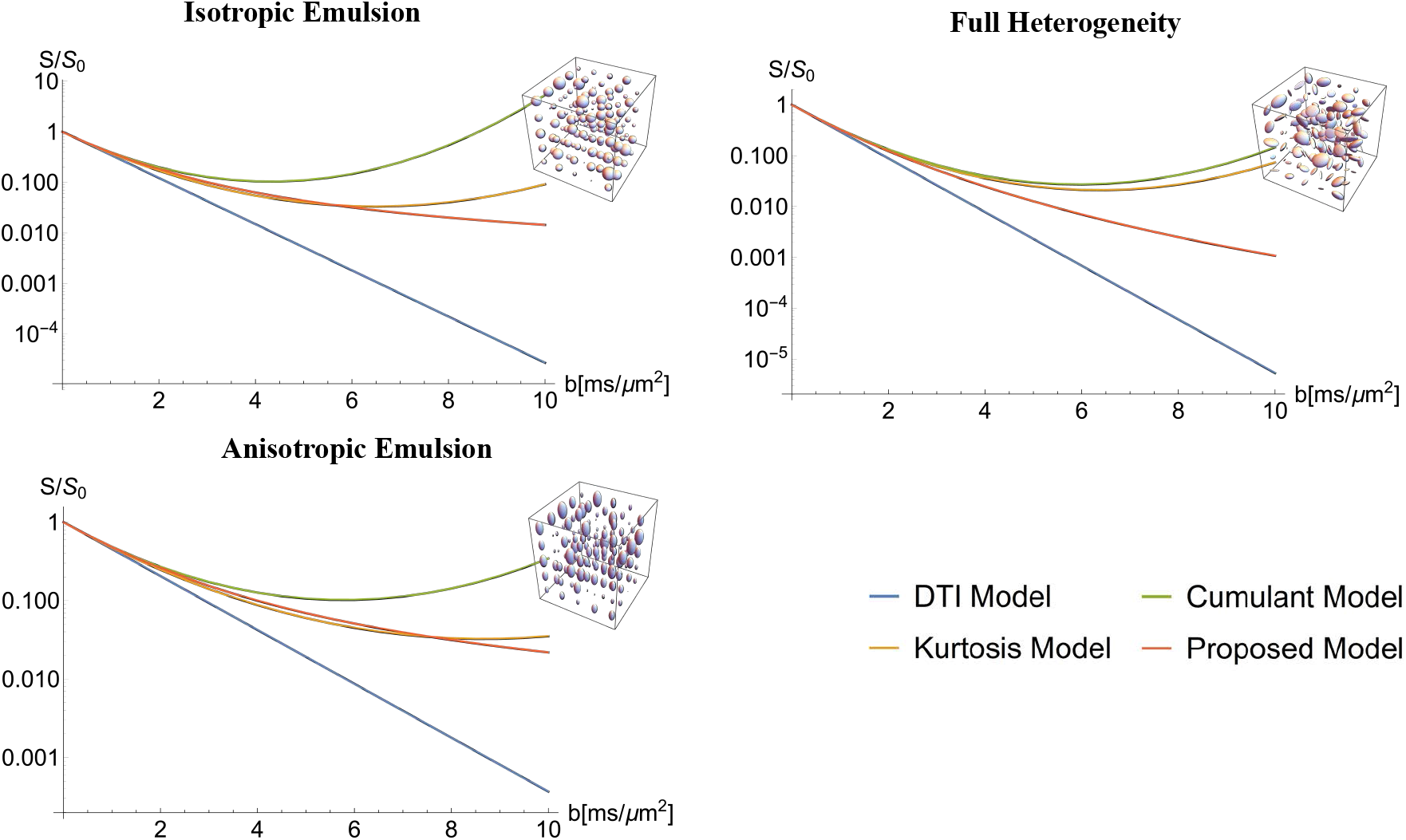
Comparison of signal models (DTI, kurtosis, cumulant and proposed model) for DTDs shown in the inset generated from CNTVD. Normalized MR signal *S*/*S*_0_ is plotted for various b-values from rank-2 b-matrices.

The well-posedness of the signal inversion problem is demonstrated in Figure 5 for various DTDs with clinically feasible b-values. The figure compares the actual signal with that calculated using the estimated parameters in the absence of noise. For our demonstration we chose DTDs with isotropic mean with size or shape heterogeneity, and anisotropic mean with orientation heterogeneity (i.e. crossing fibers). The signal estimate from the DTI model is also shown for reference. The estimated and actual mean/covariance glyphs are overlaid for comparison. The estimation framework was able to capture all the tested mean-covariance pairs with errors less than 10% likely due to convergence criterion of the optimization algorithm and/or choice of initial parameter estimate.

**Figure 5:**
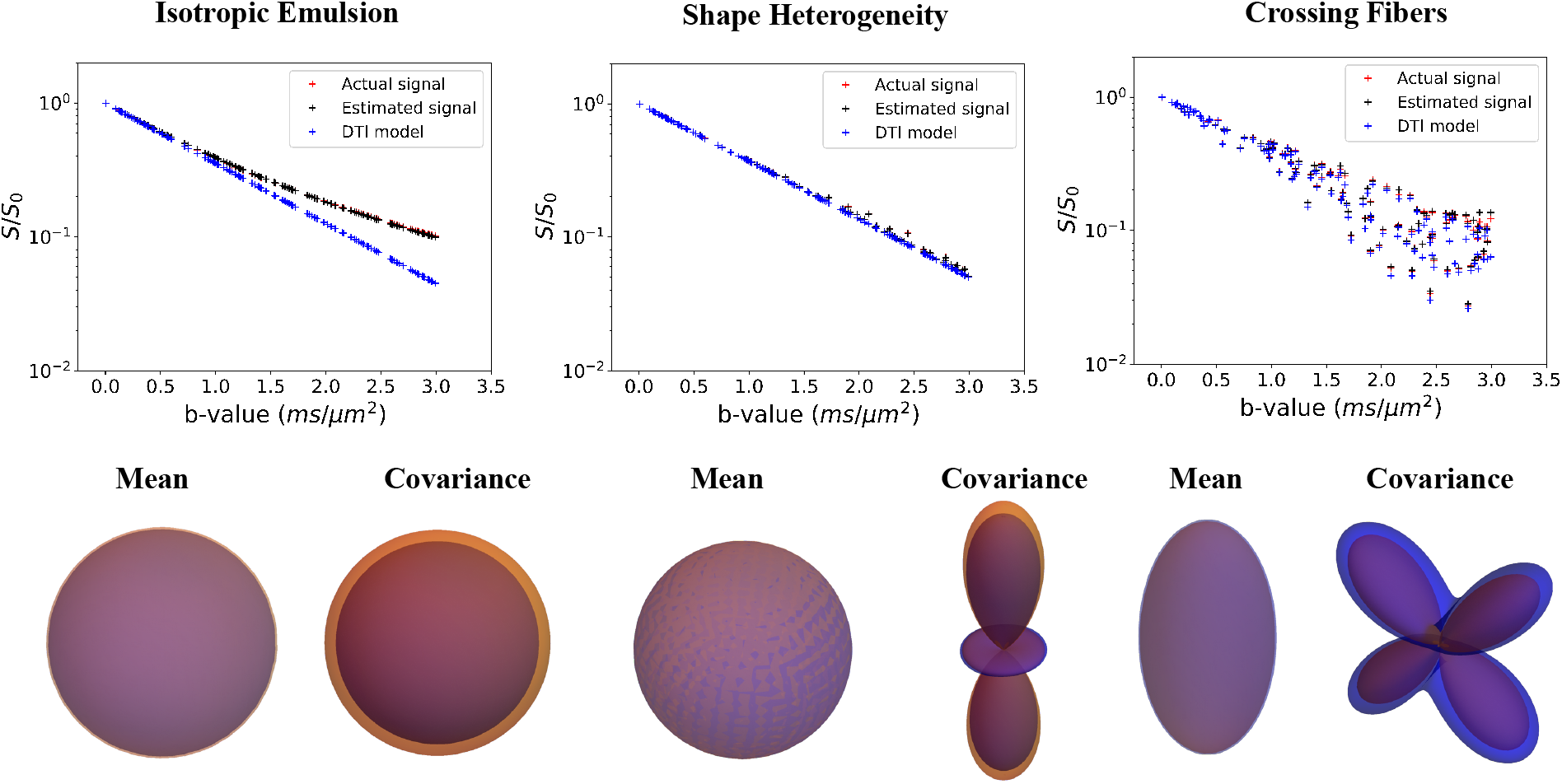
Estimation of mean and covariance tensors from synthetic MR signal without noise. The estimation is shown for three different DTDs shown in Figures 2 and 3. The signal curve from the proposed and DTI model is shown in the first row. The mean and covariance glyphs for actual (orange) and estimated (blue) DTD are overlaid and shown in the second row.

The robustness of the estimation pipeline to noise in the MR signal is shown in Figure 6. Three different realistic signal-to-noise ratios (SNR at *b* = 0 ms/*μ*m^2^) are tested for isotropic emulsion type DTD. The estimated and actual mean/covariance glyphs are overlaid for comparison similar to the previous figure. The mean tensor shape and size was exactly captured for all the SNRs. The structure of the covariance was also estimated accurately for all the three SNRs with slight error in the size likely due to reasons discussed in the previous paragraph.

**Figure 6:**
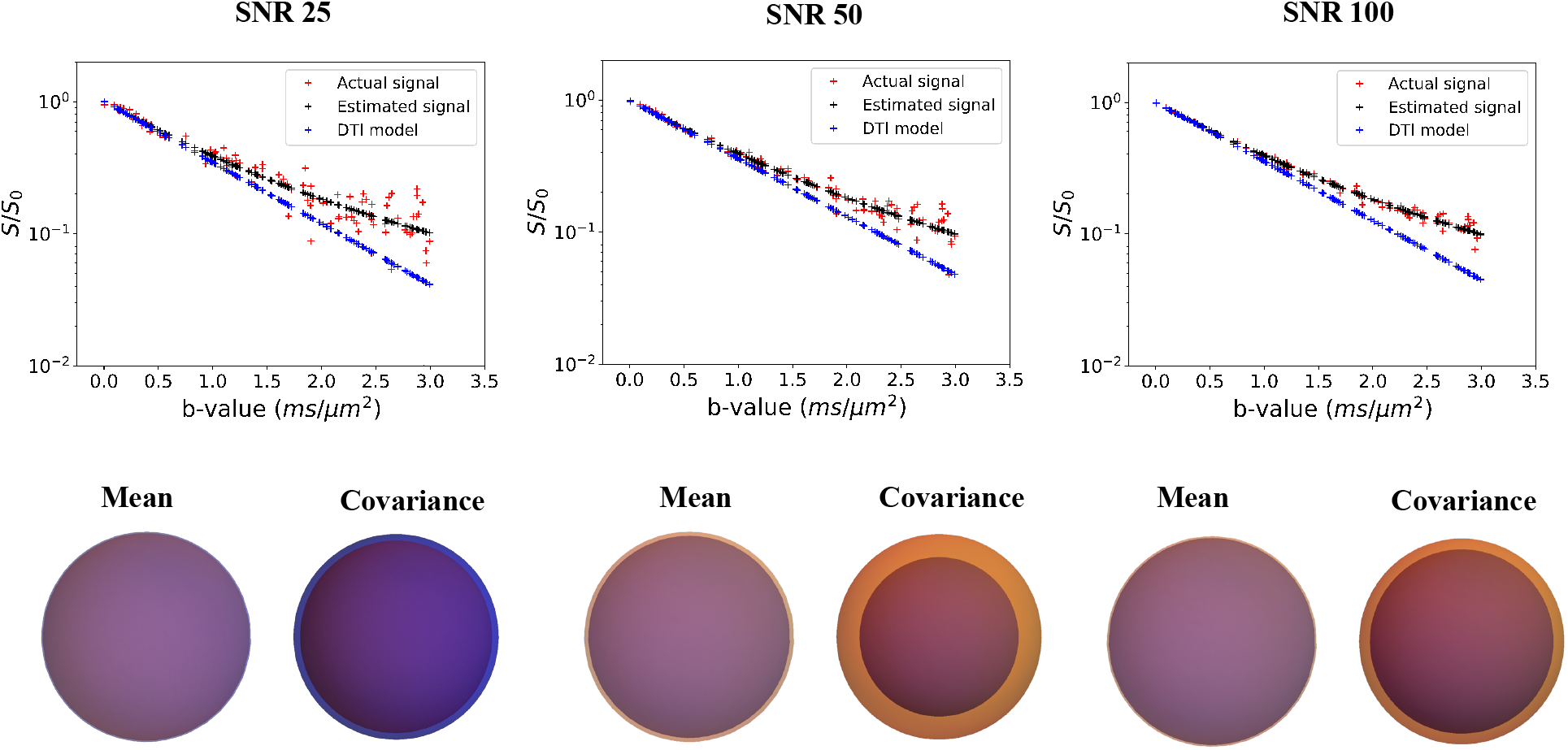
Estimation of mean and covariance tensors from synthetic MR signal for isotropic emulsion type DTD with varying signal-to-noise ratio (SNR). The signal curve from the proposed and DTI model is shown in the first row. The mean and covariance glyphs for actual (orange) and estimated (blue) DTD are overlaid and shown in the second row.

The various DTDs are visualized using glyphs and microstructural stains in Figures 7, 8 and 9. The glyphs for gray matter DTDs are shown in Figure 7. The uniform DTD has an isotropic mean tensor, zero covariance and isotropic macro and micro ODFs as expected. Size and fully heterogeneous DTDs had the same glyphs as the uniform DTD except for the covariance, which was nonzero and isotropic. The shape heterogeneous DTD had a covariance structure peaked along the longitudinal axis because the large change in the eigenvalues occur along that direction with eigenvectors fixed. There was also a slight difference between macro and *μ*ODFs.

**Figure 7:**
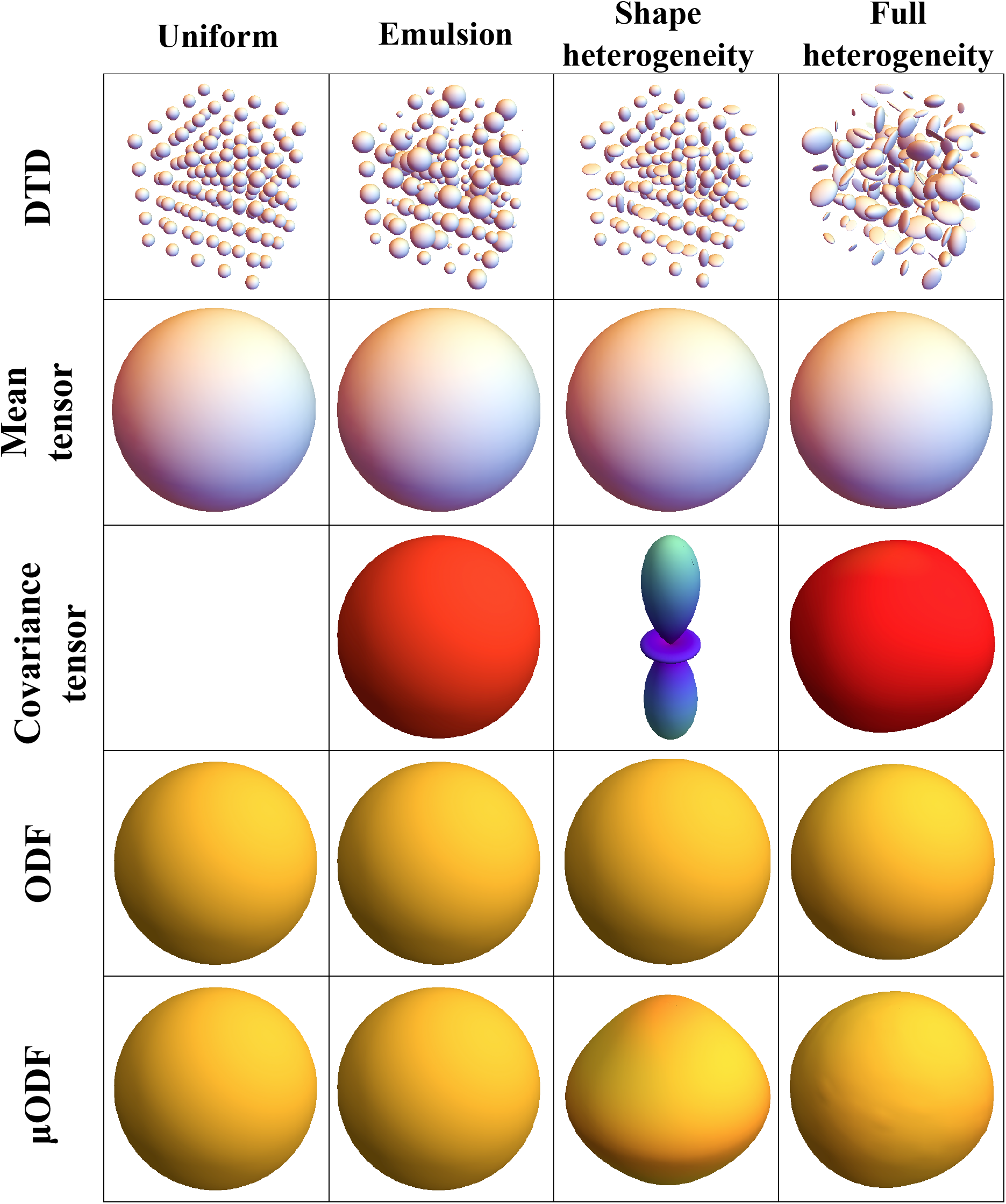
Glyphs describing macroscopically isotropic and microscopically heterogeneous DTDs generated from CNTVD. (First row) DTDs describing uniform, size, shape and orientation/size/shape heterogeneous voxels respectively. (Second row) Second order mean tensor displayed as an isosurface for each of the DTDs. (Third row) Fourth order covariance tensor projected in 3D for each of the DTDs. (Fourth row) Macro ODF obtained from the mean diffusion tensor. (Fifth row) Micro ODF obtained by averaging the ODFs of individual micro diffusion tensors in the voxel.

**Figure 8:**
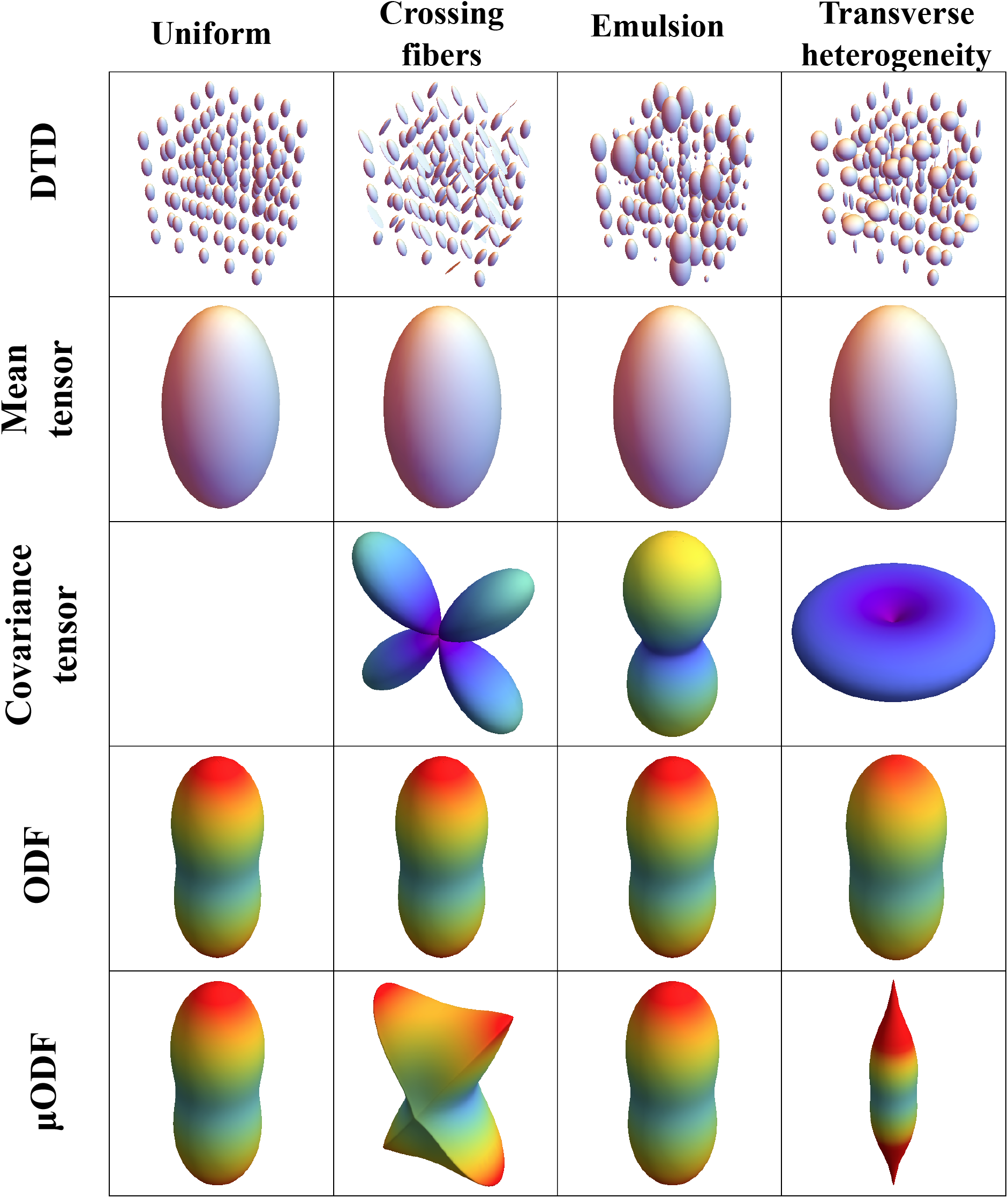
Glyphs describing macroscopically anisotropic and microscopically heterogeneous DTDs generated from CNTVD. (First row) DTDs describing uniform, crossing fibers, emulsion and transverse size heterogeneous voxels respectively. (Second row) Second order mean tensor displayed as an iso-surface for each of the DTDs. (Third row) Fourth order covariance tensor projected in 3D for each of the DTDs. (Fourth row) Macro ODF obtained from the mean diffusion tensor. (Fifth row) Micro ODF obtained by averaging the ODFs of individual micro diffusion tensors in the voxel.

**Figure 9:**
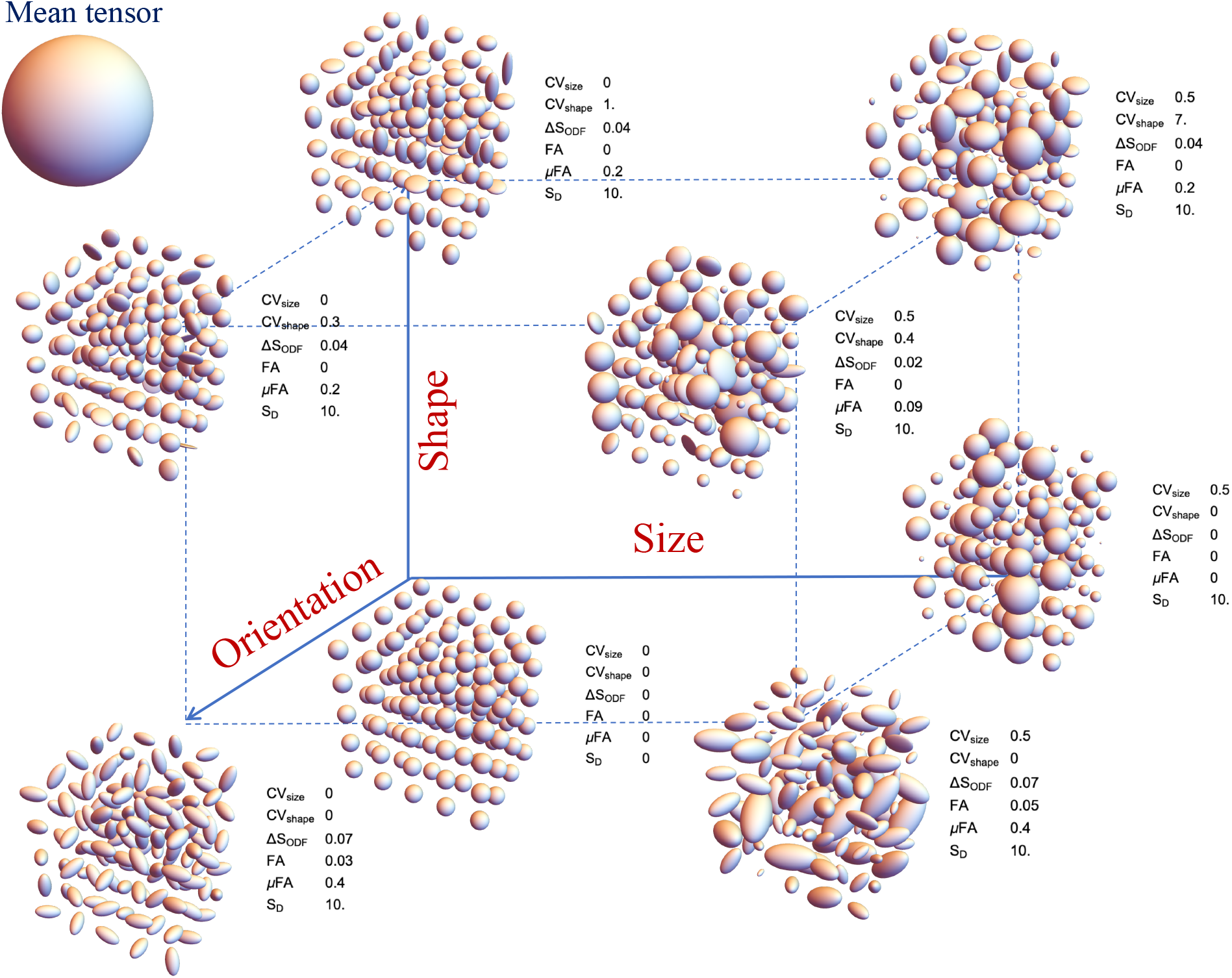
Various parametric stains depicting the shape, size and orientation heterogeneity for a macroscopically isotropic mean tensor shown on the upper left corner. The DTD and calculated values of the various stains are shown at various points in the 3D grid. Size axis refers to heterogeneity in the trace of the diffusion tensor, shape axis refers to extent of prolate-oblate-spherical mixture in the DTD, and the orientation axis refers to the dispersion in the principal direction of the diffusion tensors. CV_size_ - coefficient of variation for trace of ADC, CV_shape_ - coefficient of variation for ratios of diffusion tensor eigenvalues (i.e. *λ*_1_/*λ*_2_ and *λ*_2_/*λ*_3_), ΔS_ODF_ - ODF entropy gained due to averaging, FA - fractional anisotropy, *μ*FA - Microscopic fractional anisotropy, and S_D_ - overall entropy of the DTD.

The glyphs for white matter DTDs are shown in Figure 8. The mean tensor was prolate shaped, and ODF was peanut shaped for all the DTDs considered. The structure of the covariance and *μ*ODF was however unique for each DTD. The crossing fiber covariance and *μ*ODF had four lobes aligned with the two fiber populations. The covariance and *μ*ODF for emulsion type DTD had two lobes along the principal direction of the mean diffusion tensor. The covariance for the multi diameter fiber bundle was pancake shaped showing the heterogeneity in the transverse plane. The *μ*ODF was pill shaped due to hindered diffusion heterogeneity in the transverse plane compared to the longitudinal axis.

The various microstructural stains calculated for a variety of DTDs are shown graphically in Figure 9. The size heterogeneity metric, CV_size_, is zero for shape and orientation heterogeneous DTD but non-zero for size heterogeneous DTD. The shape metric, CV_shape_, is zero for size and orientation heterogeneous DTDs but non-zero for shape heterogeneous DTD. Thus the size and shape metrics independently capture the specific heterogeneity as desired. The orientation metric is zero for size heterogeneous DTD but non-zero for shape and orientation heterogeneous DTD. The *μ*FA was greater than FA only for shape and/or orientation heterogeneous DTD hence it fails to capture the size heterogeneity. The entropy measure was overall higher for size, shape and orientation heterogeneous DTDs compared to the uniform DTD.

## Discussion

### CNTVD for DTD

The various white and gray matter DTDs demonstrate the richness of CNTVD in describing a variety of realistic sub-domains that may exist in a voxel in the brain. Given the trace of the micro-diffusion tensors are similar, these simulated DTDs also show that CNTVD could capture multiple types of heterogeneity that maybe present within a MRI voxel. However, it should be noted that the DTD in a voxel is generally unknown, and CNTVD is proposed as one of the admissible candidate distributions for its biological resemblance in brain tissue along with useful properties such as well-posed signal inversion, maximum entropy and producing positive definite diffusion tensors. Despite the richness of the CNTVD, it however does not span the entire space of positive semi-definite diffusion tensors.

CNTVD for example could not describe situations with only orientational heterogeneity (i.e., a powder pattern) which is known to have DTD moments greater than two [39]. However, the presence of such powder patterns in brain tissue is debatable given the experimental evidence for Gaussian DTD for the orientationally averaged diffusivity [23]. Simulations showed that CNTVD could also not capture crossing fibers with angular separations not equal to 90 degrees which require a mixture type model. It should be noted that the presented framework could be applied to any DTD not restricted to CNTVD.

### Signal model

The signal curves show the effectiveness of the DTI model fitting the signal attenuation data at low b-values and the need for large b-value data to observe the effects of intravoxel heterogeneity. The large b-values also sensitizes the signal to multiple DTD compartments such as intra- and extra-cellular environments which can be captured by expressing the proposed signal equation as a weighted sum of multiple DTDs. With the above modification, the limiting b-value for our proposed model is determined by the extent of diffusion independent signal loss such as due to MR relaxation and proton density unlike the cumulant expansion and kurtosis model.

The versatility of the proposed signal model is demonstrated by its validity at large b-values excluded by the DTI, kurtosis models and cumulant representation, and for any *p*(**D**) not restricted to CNTVD. However, a central assumption of the model is the Gaussian diffusion of water within the tissue micro-compartments. A classic test for Gaussian diffusion is the time independence of diffusivity. The time-dependent diffusion in neural tissue is however experimentally observed only at very short diffusion times on the order of hundreds of microseconds [40]. The change in diffusivity in gray and white matter with diffusion times beyond a few milliseconds is very small (< 10% change from tens of milliseconds to two seconds) [41, 42, 43]. Further the breakdown of rotational invariance property of trace of the diffusion tensor, a known consequence of time-dependent diffusion [44], was also found to be very small (< 5%) in neural tissue [45]. The apparent time independent diffusion at long diffusion times typically probed in diffusion MRI experiments [46, 47] could be partly attributed to pore saturation of wide diffusion gradient pulses, water permeability of lipid membranes which hinders the diffusion rather than restricting it, and/or approach to Gaussianity due to ensemble averaging of microscopic diffusion environments within a MRI voxel. The currently available data thus seems to suggest that at time scales of 20-60 ms typically probed in standard spin echo (SE) diffusion weighted scans, the water diffusion in the brain parenchyma is approximately Gaussian.

### Experimental Design

The estimation pipeline was accurate and robust to noise which is partly attributed to the model selection framework and the novel experimental design which uniformly samples b-matrices of varying size, shape and orientation. The equivalence between the covariance matrix and fourth order covariance tensor representation introduced in [27] enabled us to exploit the symmetries of the general fourth order tensor to build a family of nested models for covariance estimation.

The importance of the choice of diffusion encoding gradient vectors in the estimation of the diffusion tensor has been well studied [48, 49]. It can be expected that the same holds true for DTD estimation as well especially for higher rank b-matrix measurements requiring multiple gradient pulses which increases the number of degrees of freedom. An optimal experimental design which significantly reduces the number of measurements is essential for practical applicability of the method. Komlosh et al. used rank-2 b-matrices obtained using orthogonal gradient pairs to extract microscopic heterogeneity in excised spinal cord [17]. Jespersen et al. introduced a *t*-design scheme for rank-2 b-matrix measurements to estimate orientation heterogeneous DTDs in the excised brain [50]. The two gradient vectors were made orthogonal to each other and chosen as the vertices of the icosahedron. The orthogonality constraint on the gradient vectors restricts the space of positive definite matrices explored, while sampling antipodal vectors leads to redundant b-matrices which ill-conditions the DTD signal inversion. Westin et al. used a multi-shell approach where each shell is composed of a number of rank 1, 2 and 3 b-matrices to account for arbitrary DTDs in the brain *in-vivo* [24]. Martins and Topgaard uniformly sampled the space of orientation, shape and size of the b-matrices in a phantom experiment which is extremely time consuming [51].

Our experimental design using CS balances acquisition time and the space of fourth order covariance tensors spanned by the b-matrices [52]. A continuum of b-matrices of varying shapes, sizes and orientations were chosen instead of a multi-shell acquisition to get a better definition of the MR signal dependence with b-value. Such an experimental design allows an unbiased estimation of the DTD parameters in a reasonable amount of time. The lack of an extra gradient pair for rank-3 b-matrices, which were shown to be not necessary to estimate the covariance, reduces the number of degrees of freedom to explore and possibly the echo time of the acquisition particularly in clinical MRI scanners which have long gradient ramp times.

### DTD visualization

The presented glyphs and stains provide an unique information about the underlying micro structure as shown in the figures. The observed difference between macro- and micro- ODFs, and non-zero value of orientation heterogeneity metric for shape heterogeneous DTDs is due to the fact that changing the shape of the diffusion tensor from prolate to oblate changes the principal axis of the diffusion tensor from longitudinal to transverse which slightly mimic as orientation heterogeneity. We also show that the *μ*FA does not fully capture the heterogeneity in the tissue thus the need for additional stains.

The proposed stains are generic and applies to any of the eight covariance models. They could help inform about normal and abnormal micro structural changes in tissue with disease. For Gaussian diffusion, the ODF glyph purely characterizes the orientation dependence of diffusivity independent of experimental parameters such as diffusion time. Comparing the macro and micro ODF glyphs could be particularly revealing in white matter voxels. The size metric could help distinguish between layers of the cerebral cortex which are known to have different cell size distribution that gets altered in diseases such as schizophrenia and Huntington’s disease [53]. It could also reveal the cytoarchitecture in spinal cord where diameter distribution of various fiber tracts and nerve roots traversing the cord vary widely [54, 55, 56].

## Conclusion

In this study, a novel paradigm to simulate and estimate DTD is presented. A new MC based signal model is introduced which is monotonically decreasing with b-value unlike previous models and designed to work for any DTD. The richness of CNTVD is shown using a set of realistic DTDs that may exist in brain gray and white matter. The data inversion is performed using a parsimonious approach to accurately estimate various mean and covariance pairs in the presence of noise in the MR signal. A family of new novel parametric stains and glyphs are shown to capture distinct features of micro structural inhomogeneity within a MRI voxel. The method is thus an important advance in the continuing quest to explore the microstructure within the MRI voxel.

## Supporting information

Supplementary Proofs

## Acknowledgments

We would like to thank Dan Benjamini for thoughtful discussions. PJB, SP, and KNM were all supported by the Intramural Research Program of the *Eunice Kennedy Shriver* National Institute of Child Health and Human Development. DG was supported under the auspices of the Center for Neurosciences and Regenerative Medicine (CNRM) and the Henry Jackson Foundation (HJF). This work was partly funded from NIH BRAIN initiative grant, 1U01EB026996-01 - Connectome 2.0: Developing the next generation human MRI scanner for bridging studies of the micro-, meso- and macro-connectome.

## Author Contributions

PJB designed the research. DG wrote the proofs and assisted with estimation. SP performed analysis on metrics and assisted speeding up the computation. MK carried out the simulation/estimation and wrote the article. SP, PJB and DG edited the article. All authors reviewed the manuscript.

