## Supplementary Proofs for "A New Framework for MR Diffusion Tensor Distribution"

#### 1 Maximum entropy property and identifiability of the CNTVD

**Theorem 1.** *For a Borel-measurable set, $\mathcal{C} \subseteq \mathbb{R}^d$, the constrained Gaussian distribution with density,*

$$p_{\mu,\Omega}(X|X \in \mathcal{C}) = \frac{1}{Z(\mu,\Omega)} \exp\left(-\frac{1}{2}(X-\mu)\Omega(X-\mu)^\top\right)\mathbf{1}(X \in \mathcal{C})$$

*where $Z(\mu,\Omega)$ is the normalizing constant is the unique maximizer of the entropy*

$$H(p) = -\int_{\mathcal{C}} p(X)\log p(X)dX$$

*among all probability densities supported by $\mathcal{C}$ with given first and second moments*

$$\langle X_k \rangle = \mathbb{E}_{\mu,\Omega}(X_k|X \in \mathcal{C}), \quad \langle X_k X_\ell \rangle = \mathbb{E}_{\mu,\Omega}(X_k X_\ell|X \in \mathcal{C}), \quad 1 \leq k, \ell \leq d.$$

*When $\mathcal{C}$ contains an open ball the parameters $\mu, \Omega$ are uniquely determined by the first and second moments.*

*Proof.* Consider the following Lagrangian with the constraint $p(X) \geq 0$,

$$\mathcal{L}(p,\lambda,\nu,\Xi) = -\int_{\mathcal{C}} p(X)\log p(X)dX + \left(1 - \int_{\mathcal{C}} p(X)dX\right)\lambda + \left(\langle X \rangle - \int_{\mathcal{C}} Xp(X)dX\right)\cdot \nu \quad (1)$$

$$+ \text{Trace}\left[\left(\langle X^\top X \rangle - \int_{\mathcal{C}} X^\top Xp(X)dX\right)\Xi\right] \quad (2)$$

where $\lambda \in \mathbb{R}, \nu \in \mathbb{R}^d$ and $\Xi \in \mathbb{R}^{d\times d}_{symm}$ are the Lagrange multipliers. A solution of the maximization problem corresponds necessarily to a zero of the variational derivatives,

$$\frac{\delta\mathcal{L}}{\delta p(X)} = -\log p(X) - 1 + \lambda + X\cdot\nu + X\Xi X^\top = 0 \quad \forall X \in \mathcal{C} \tag{3}$$

which means that when the maximum entropy solution exists necessarily it has the form

$$p(X) = \exp\Big(\lambda - 1 + X\cdot\nu + X\Xi X^\top\Big)\mathbf{1}(X \in \mathcal{C}) \tag{4}$$

with $\lambda$ corresponding to a normalizing constant and $\nu, \Xi$ such that the first and second moment constraints are satisfied. When a solution with the given moments exists, after reparametrization, the density is rewritten as

$$p(X) = \frac{1}{Z(\mu,\Omega)}\exp\Big(-\frac{1}{2}(X-\mu)\Omega(X-\mu)^\top\Big)\mathbf{1}(X \in \mathcal{C}). \tag{5}$$

When the support $\mathcal{C}$ is unbounded, necessarily $\Omega$ is positive definite on the recession cone of $\mathcal{C}$, and when $\Omega$ is positive definite on $\mathbb{R}^d$, we obtain a constrained Gaussian distribution. When a maximum exists, the uniqueness of the maximizing parameters is shown below.

Suppose that $p_{\widetilde{\mu},\widetilde{\Omega}}(X)$ is another constrained Gaussian distribution with support $\mathcal{C}$ satisfying the moment constraints and maximizing the entropy. Then since the distributions satisfy the same moment constraints,

$$\nabla \log Z(\mu,\Omega) = \nabla \log Z(\widetilde{\mu},\widetilde{\Omega}) = \big(\langle X\rangle, \langle X^\top X\rangle\big) \tag{6}$$

which implies

$$\begin{aligned} 0 &= \big(\nabla \log Z(\mu,\Omega) - \nabla \log Z(\widetilde{\mu},\widetilde{\Omega})\big)\big(\mu-\widetilde{\mu}, \Omega-\widetilde{\Omega}\big)^\top \quad (7)\\ &= \big(\mu-\widetilde{\mu}, \Omega-\widetilde{\Omega}\big)\ \nabla^\top\nabla \log Z(\hat{\mu},\hat{\Omega})\ \big(\mu-\widetilde{\mu}, \Omega-\widetilde{\Omega}\big)^\top \quad (8)\end{aligned}$$

with $\hat{\mu} = \alpha\mu + (1-\alpha)\widetilde{\mu}$ and $\hat{\Omega} = \alpha\Omega + (1-\alpha)\widetilde{\Omega}$ for some $0 \le \alpha \le 1$, and

$$\begin{aligned} \nabla^\top\nabla \log Z(\hat{\mu},\hat{\Omega}) &= \frac{\nabla^\top\nabla Z(\hat{\mu},\hat{\Omega})}{Z(\hat{\mu},\hat{\Omega})} - \frac{\nabla^\top Z(\hat{\mu},\hat{\Omega})\nabla Z(\hat{\mu},\hat{\Omega})}{Z(\hat{\mu},\hat{\Omega})^2} \quad (9)\\ &= \begin{bmatrix} \mathrm{Cov}_{\hat{\mu},\hat{\Omega}}(X) & \mathrm{Cov}_{\hat{\mu},\hat{\Omega}}(X^\top X, X) \\ \mathrm{Cov}_{\hat{\mu},\hat{\Omega}}(X, X^\top X) & \mathrm{Cov}_{\hat{\mu},\hat{\Omega}}(X^\top X, X^\top X) \end{bmatrix} \quad (10)\end{aligned}$$

which means that

$$\mathrm{Variance}_{\hat{\Omega},\hat{\theta}}\Big((\mu-\widetilde{\mu})X^\top + X(\Omega-\widetilde{\Omega})X^\top\Big) = 0. \tag{11}$$

Since $p_{\hat{\mu},\hat{\Omega}}(X) > 0$ on $\mathcal{C}$,

$$\mathcal{C} \subseteq \left\{X \in \mathbb{R}^d : (\mu - \widetilde{\mu})X^\top + X(\Omega - \widetilde{\Omega})X^\top = r\right\} \tag{12}$$

for some $r \in \mathbb{R}$, which is a variety of dimension, $(d-1)$, when $\mu \neq \widetilde{\mu}$ or $\Omega \neq \widetilde{\Omega}$, contradicting the assumption of $\mathcal{C}$ having full dimension □

### 2 Relations between the diffusion tensor cumulants and the displacement cumulants

Let $X \in \mathbb{R}^d$ be a conditionally Gaussian displacement vector, with zero conditional mean and conditional covariance $2D$, while $D$ is a random symmetric positive definite matrix (diffusion tensor) with probability density $p(D)$. The Characteristic Function (CF) of the Gaussian conditional law of $X$ given $D$ is given by

$$\mathbb{E}\big(\exp(iq \cdot X)\big|D\big) = \exp\big(-qDq^\top\big) \tag{13}$$

while the unconditional CF is obtained by integrating w.r.t. $p(D)$ the conditional CF

$$\mathbb{E}\big(\exp(iq \cdot X)\big) = \int_{\mathcal{M}_+} \exp\big(-qDq^\top\big)p(D)dD = \mathbb{E}\big(\exp(-\mathrm{Trace}(QD))\big), \quad q \in \mathbb{R}^d, \tag{14}$$

and it coincides with the Laplace Transform (LT) of the diffusion tensor distribution $p(D)$ at $Q = q^\top q$.

We introduce the following notations: the order of a multi-index $\alpha = (\alpha_1, \ldots, \alpha_d) \in \mathbb{N}^d$ is denoted by

$$|\alpha| = \sum_{j=1}^{d} \alpha_j \tag{15}$$

and $q^\alpha = \prod_{j=1}^{d} q_j^{\alpha_j}$ for $q \in \mathbb{R}^d$. The multivariate cumulants of $X = (X_1, \ldots, X_d)$ are denoted by

$$\kappa_\alpha(X) = \kappa_{|\alpha|}\big(\underbrace{X_1, \ldots, X_1}_{\alpha_1 \text{ copies}}, \underbrace{X_2, \ldots, X_2}_{\alpha_2 \text{ copies}}, \ldots, \underbrace{X_d, \ldots, X_d}_{\alpha_d \text{ copies}}\big), \tag{16}$$

so that $\kappa_{e_j+e_\ell}(X) = \kappa_2(X_j, X_\ell)$, $\kappa_{e_j+e_k+e_\ell+e_r}(X) = \kappa_4(X_j, X_k, X_\ell, X_r)$, where $\{e_1, \ldots, e_d\}$ is the cartesian basis.

The cumulant expansion of the CF of $X$ is given by

$$\log \mathbb{E}\big(\exp(iq\cdot X)\big) = \sum_{\alpha:|\alpha|\in 2\mathbb{N}\setminus\{0\}} \frac{\kappa_\alpha(X)}{\alpha!}(-1)^{|\alpha|/2} q^\alpha \tag{17}$$

$$= \sum_{n=1}^{\infty} \frac{(-1)^n}{(2n)!} \sum_{j_1\dots j_{2n}=1}^{d} \kappa_{2n}(X_{j_1},\dots,X_{j_{2n}}) q_{j_1}\dots q_{j_{2n}} \tag{18}$$

where all the odd moments and cumulants vanish since the law of $X$ is reflection symmetric, while the cumulant expansion of the LT of $D$ is given by

$$\log \mathbb{E}\big(\exp(-\mathrm{Trace}(QD))\big) = \sum_{\beta:|\beta|>0} \frac{\kappa_\beta(D)}{\beta!}(-1)^{|\beta|} 2^{|\bar{\beta}|} Q^\beta \tag{19}$$

$$= \sum_{n=1}^{\infty} \frac{(-1)^n}{n!} \sum_{j_1\dots j_{2n}=1}^{d} \kappa_n(D_{j_1 j_2},\dots,D_{j_{2n-1}j_{2n}}) Q_{j_1 j_2}\dots Q_{j_{2n-1}j_{2n}} \tag{20}$$

where $\beta = (\beta_{ij} : 1 \le i \le j \le d) \in \mathbb{N}^{(d+1)d/2}$, and $\bar{\beta}$ is the restriction of the multi-index $\beta$ to the off-diagonal entries. By comparing these expansions for rank-1 $Q = q^\top q$, we see that

$$\langle X\rangle = 0, \quad \langle X^\top X\rangle = \mathrm{Cov}(X) = 2\langle D\rangle, \tag{21}$$

$$\kappa_4(X_i,X_j,X_k,X_\ell) = 4\big\{\mathrm{Cov}(D_{ij},D_{k\ell}) + \mathrm{Cov}(D_{ik},D_{j\ell}) + \mathrm{Cov}(D_{i\ell},D_{jk})\big\} \tag{22}$$

$$\kappa_{2n}(X_{j_1},\dots,X_{j_{2n}}) = 2^n \sum_{\pi=\{\pi_1,\dots,\pi_n\}} \kappa_n(D_{\pi_1},\dots,D_{\pi_n}) \tag{23}$$

where the sum in Equation (23) is over the pairings $\pi$ of $\{j_1,\dots,j_{2n}\}$ into pairs $\pi_1,\dots,\pi_n$, in accordance with the law of total cumulance. The even cumulants of the displacement $\kappa_\alpha(X)$ with $|\alpha| = 2n$ are, up to rescaling, symmetrizations of corresponding cumulants of the diffusion tensor $\kappa_\beta(D)$ with $|\beta| = n$ .

By truncating the cumulant expansion Equation (19) of the LT of $p(D)$ up to second order terms, we obtain the LT of the NTVD with mean $\langle D\rangle$ and Covariance$(D_{ij},D_{k\ell}) = \mathsf{C}_{ijk\ell}$, which can be used to model the signal decay in a multiple pulsed gradient experiment:

$$\frac{S(\mathsf{Q})}{S(0)} = \mathbb{E}\big[\exp\big(-\mathrm{Trace}(\mathsf{QD})\big)\big] = \exp\left(-\mathrm{Trace}(\mathsf{Q}\langle D\rangle) + \frac{1}{2}\sum_{ijk\ell} \mathsf{C}_{ijk\ell}\mathsf{Q}_{ij}\mathsf{Q}_{k\ell}\right) \tag{24}$$

For single pulsed gradient experiments $\mathsf{Q} = q^\top q$ has rank 1, and Equation (24) coincides with the signal decay in Diffusion Kurtosis Imaging (DKI) [1], obtained by truncating the cumulant expansion of the CF of $X$ (Equation (17)) up to 4th order terms,

$$\frac{S(q)}{S(0)} = \exp\left(-q\overline{\mathsf{D}}q^\top + \frac{1}{6}\mathrm{Trace}(\overline{\mathsf{D}})^2 \sum_{ijk\ell} \mathsf{K}_{ijk\ell} q_i q_j q_k q_\ell\right), \tag{25}$$

where

$$\overline{\mathsf{D}} = \langle \mathsf{D} \rangle \quad \text{and } \mathsf{K}_{ijk\ell} = \frac{\mathsf{C}_{ijk\ell} + \mathsf{C}_{ikj\ell} + \mathsf{C}_{i\ell jk}}{\text{Trace}(\langle \mathsf{D} \rangle)^2}. \tag{26}$$

are respectively the diffusion tensor and the diffusion kurtosis tensor. Thus the NTVD signal model extends the DKI signal model from single to multiple pulsed experiments, and up to rescaling, the diffusion kurtosis tensor is the symmetrization of the partially symmetric NTVD covariance tensor. Both NTVD and DKI signal models are "non-physical" and break down at high $b$-value, since the NTVD supports also tensors which are not positive definite, and the DKI signal decay is not the CF of a probability distribution.

### 3 Sufficiency of rank-1 and 2 b-matrices for diffusion tensor covariance estimation

**Lemma 1.** *The linear space $\mathcal{T}$ of partially symmetric 4th-order d-dimensional tensors with symmetries*

$$\mathsf{T}_{jik\ell} = \mathsf{T}_{ijk\ell} = \mathsf{T}_{k\ell ij} \quad \forall 1 \le i, j, k, \ell \le d$$

*is spanned by tensors of the form*

$$(u \otimes u + v \otimes v) \otimes (u \otimes u + v \otimes v) \quad \textit{with } u, v \in \mathbb{R}^d,$$

*where $\otimes$ denotes outer product, and by symmetrizing it follows that the tensors $u \otimes u \otimes u \otimes u$ with $u \in \mathbb{R}^d$ span the subspace of totally symmetric 4th-order tensors.*

*Proof.* By the spectral decomposition, every $\mathsf{T} \in \mathcal{T}$ is a linear combination of at most $\frac{(d+1)d}{2}$ 4th-order tensors of the form

$$\left(\sum_{k=1}^{d} u_k \otimes u_k\right) \otimes \left(\sum_{\ell=1}^{d} u_\ell \otimes u_\ell\right) = \sum_{k,\ell=1}^{d} u_k \otimes u_k \otimes u_\ell \otimes u_\ell \tag{27}$$

$$= \frac{1}{2} \sum_{k,\ell=1}^{d} (u_k \otimes u_k + u_\ell \otimes u_\ell) \otimes (u_k \otimes u_k + u_\ell \otimes u_\ell) - d \sum_{k=1}^{d} u_k \otimes u_k \otimes u_k \otimes u_k \tag{28}$$

with $u_k \in \mathbb{R}^d$, where

$$u \otimes u \otimes v \otimes v + v \otimes v \otimes u \otimes u = (u \otimes u + v \otimes v) \otimes (u \otimes u + v \otimes v) - u \otimes u \otimes u \otimes u - v \otimes v \otimes v \otimes v \tag{29}$$

□

It can be observed that Equation (28) is a sum of outer squares of rank-1 and rank-2 b-matrices.

Thus, rank-3 b-matrices are not necessary and rank-1 b-matrices are not enough to estimate the covariance of the diffusion tensor in a multiple pulsed gradient experiment.
